# All thresholds of maternal hyperglycaemia from the WHO 2013 criteria for gestational diabetes identify women with a higher genetic risk for type 2 diabetes

**DOI:** 10.1101/671057

**Authors:** Alice E Hughes, M. Geoffrey Hayes, Aoife M Egan, Kashyap A Patel, Denise M Scholtens, Lynn P Lowe, William L Lowe, Fidelma P Dunne, Andrew T Hattersley, Rachel M Freathy

## Abstract

**Background:** Using genetic scores for fasting plasma glucose (FPG GS) and type 2 diabetes (T2D GS), we investigated whether the fasting, 1-hour and 2-hour glucose thresholds from the WHO 2013 criteria for gestational diabetes (GDM) have different implications for genetic susceptibility to raised fasting glucose and type 2 diabetes in women from the Hyperglycemia and Adverse Pregnancy Outcome (HAPO) and Atlantic Diabetes in Pregnancy (DIP) studies.

**Methods:** Cases were divided into three subgroups: (i) FPG ≥5.1 mmol/L only, n=222; (ii) 1-hour glucose post 75 g oral glucose load ≥10 mmol/L only, n=154 (iii) 2-hour glucose ≥8.5 mmol/L only, n=73); and (iv) both FPG ≥5.1 mmol/L and either of a 1-hour glucose ≥10 mmol/L or 2-hour glucose ≥8.5 mmol/L, n=172. We compared the FPG and T2D GS of these groups with controls (n=3,091) in HAPO and DIP separately.

**Results:** In HAPO and DIP, the mean FPG GS in women with a FPG ≥5.1 mmol/L, either on its own or with 1-hour glucose ≥10 mmol/L or 2-hour glucose ≥8.5 mmol/L, was higher than controls (all *P* <0.01). Mean T2D GS in women with a raised FPG alone or with either a raised 1-hour or 2-hour glucose was higher than controls (all *P* <0.05). GDM defined by 1-hour or 2-hour hyperglycaemia only was also associated with a higher T2D GS than controls (all *P* <0.05).

**Conclusions:** The different diagnostic categories that are part of the WHO 2013 criteria for GDM identify women with a genetic predisposition to type 2 diabetes as well as a risk for adverse pregnancy outcomes.

## INTRODUCTION

Gestational diabetes mellitus (GDM) has been variably defined since criteria were first developed over 50 years ago [1]. The World Health Organization (WHO) introduced diagnostic criteria for GDM in 1999, based on criteria for overt diabetes in the general population, with a fasting plasma glucose (FPG) ≥7.0 mmol/L or impaired glucose tolerance with a 2-hour glucose post 75 g oral glucose tolerance test (OGTT) ≥7.8 mmol/L, measured between 24 and 28 weeks gestation [2]. However, lesser degrees of maternal fasting hyperglycaemia have long been associated with a higher risk for adverse perinatal outcomes [3], so a FPG ≥6.1 mmol/L (indicative of impaired fasting glycaemia in the non-pregnant population [4]) was also integrated into the WHO criteria.

The Hyperglycemia and Adverse Pregnancy Outcome (HAPO) Study [5] followed 23,316 women who underwent a 2-hour OGTT between 24 and 32 weeks gestation throughout pregnancy and found a continuous association between maternal glucose values and adverse perinatal outcomes, including birth weight ≥90^th^ centile (large for gestational age, LGA) and primary caesarean section. In 2010, the International Association of Diabetes and Pregnancy Study Groups (IADPSG) determined cut-off values equivalent to 1.75 times the odds for adverse pregnancy outcomes at mean glucose values, resulting in diagnostic thresholds for FPG ≥5.1 mmol/L, 1-hour glucose ≥10 mmol/L and 2-hour glucose ≥8.5 mmol/L [6].

WHO adopted the recommendations of IADPSG in 2013 [2], which has resulted in a higher number of cases identified as GDM due to the lower FPG threshold (estimated up to 17.8% prevalence of GDM for IADPSG 2010 criteria [6] vs 9.4% prevalence for WHO 1999 criteria [7]). Whilst these thresholds were chosen for their Obstetric risks, the HAPO Follow-Up Study found that women diagnosed by the newer criteria have a higher risk of developing disorders of glucose metabolism, including T2D, 10 years after the episode of GDM [8]. A proportion of this risk can be attributed to genetic predisposition, since genome wide association study (GWAS) data from large, non-pregnant population-based studies have identified multiple loci associated with FPG [9] and type 2 diabetes [10] and some of these are shared with GDM [11–15]. Specific to the WHO 2013 criteria, single nucleotide polymorphisms (SNPs) at the *GCK* and *TCF7L2* loci were shown to be associated with FPG and 2-hour glucose levels post-OGTT in women with GDM [16]. In addition, genetic risk scores for glycaemic traits, including FPG and type 2 diabetes, have been associated with a higher odds for GDM according to the WHO 2013 criteria [17]. However, it is not known whether the underlying genetic predisposition to fasting hyperglycaemia and type 2 diabetes varies depending on how the diagnosis of GDM is met.

We used a genetic score (GS) for FPG (FPG GS) or T2D (T2D GS) (consisting of previously-identified loci [9,18]) to test the hypothesis that there may be different genetic risks for fasting hyperglycaemia and type 2 diabetes depending on the different measurements of glucose tolerance used to diagnose GDM.

## METHODS

### Study population

Women of European ancestry with singleton pregnancies and without known pre-existing diabetes from the Hyperglycemia and Adverse Pregnancy Outcome (HAPO) Study [5] (*n*=2,628) and Atlantic Diabetes in Pregnancy (DIP) study [19] (*n*=1,084) were included. The HAPO study was an observational, multi-centre study (*N*=23,316 participants from 15 centres) to which women were recruited during pregnancy if they were over 18 years of age [5]. The 2,665 European-ancestry participants included in the current study were those with genotype data available on selected SNPs (see below). The DIP study had a case-control design: approximately three genotyped control participants without GDM (defined initially as a maternal FPG <5.6 mmol/L and/or 2-hour glucose post oral glucose load <7.8 mmol/L) were available for every genotyped case participant included in our analyses. Women who were unblinded due to being diagnosed with diabetes or GDM by pre-existing criteria used at the time of the studies were not excluded from this analysis.

### Sample collection and clinical characteristics

The study methods used in HAPO and DIP have been described in detail previously [5,7,19–21]. Maternal FPG in mmol/L was measured prior to a standard 2-hour OGTT with 75 g of glucose between 24 and 32 weeks in HAPO and 24 and 28 weeks in DIP. Information on maternal age, pre-pregnancy body mass index (BMI) and systolic blood pressure (SBP, in mmHg) was collected at the OGTT appointment. Clinical characteristics of participants in HAPO and DIP with and without GDM were different (women in DIP were older, had a higher BMI and higher SBP, all *P* <0.01), hence clinical characteristics (where available) have been presented separately.

### GDM diagnostic criteria subgroups

We used the WHO 2013 cut-offs (previously IADPSG 2010) to define fasting and 2-hour hyperglycaemia. Thus, in the current study, women diagnosed with GDM were divided into fasting hyperglycaemia only (FPG ≥5.1 mmol/L and 1-hour and 2-hour glucose post 75 g oral glucose load <10 mmol/L and <8.5 mmol/L, respectively, n=222), elevated 1-hour glucose only (1-hour glucose ≥10 mmol/l, FPG <5.1 mmol/L and 2-hour glucose <8.5 mmol/l, n=154), elevated 2-hour glucose only (2-hour glucose ≥8.5 mmol/L, FPG <5.1 mmol/L and 1-hour glucose <10 mmol/L, n=73) and both (FPG ≥5.1 mmol/L and either a 1-hour glucose ≥10 mmol/L or 2-hour glucose ≥8.5 mmol/L, or both, n=172) subgroups (Figure 1). Women without GDM were defined as having FPG <5.1 mmol/L, 1-hour glucose <10 mmol/L and 2-hour glucose <7.8 mmol/L (n=3,091). The distributions of the women in the different groups and in each of the study cohorts are shown in Figure 1.

**Figure 1.**
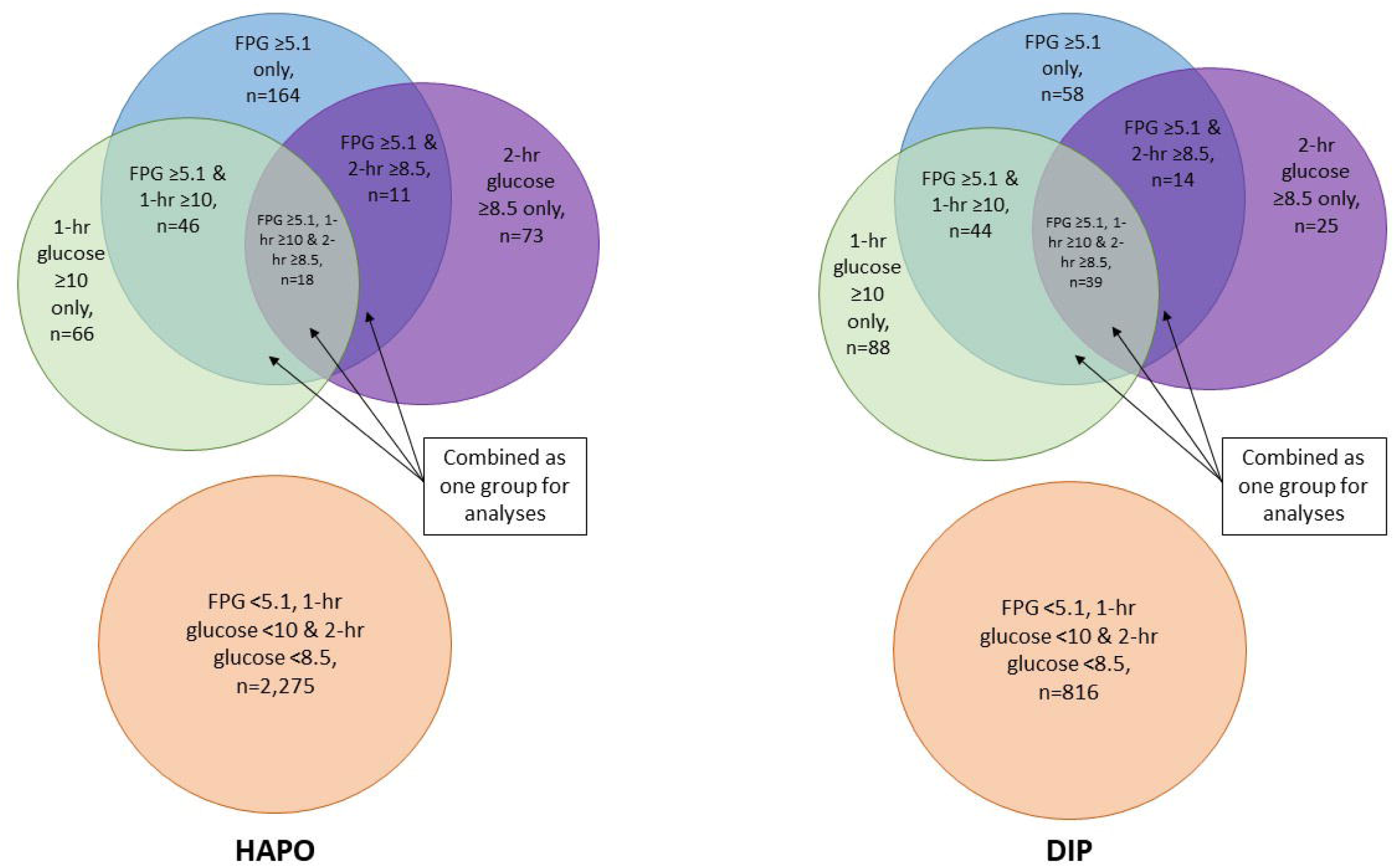
Distribution of participants diagnosed with GDM by different glucose categories in HAPO and DIP. All glucose values are in mmol/L. The 1-hr and 2-hr glucose measures refer to the glucose level measured at 1 and 2 hours, respectively, following a 75 g oral glucose load as part of an oral glucose tolerance test. Women with a FPG ≥5.1 mmol/L and either a 1-hr glucose ≥10 mmol/L or 2-hr glucose ≥8.5 mmol/L, or both, were combined as one group for analyses. DIP; Atlantic Diabetes in Pregnancy; FPG, fasting plasma glucose; GDM, gestational diabetes; HAPO; Hyperglycemia and Adverse Pregnancy Outcome Study.

### Genotyping

Genotyping of individual SNPs in DNA samples from both the DIP and HAPO studies was carried out at LGC Genomics (Hoddesdon, UK; https://www.lgcgroup.com), using the PCR-based KASP™ genotyping assay. We first selected 41 SNPs that had been previously associated with type 2 diabetes, and 16 SNPs associated with fasting glucose in non-pregnant individuals, for genotyping in the DIP study. Overlap between the type 2 diabetes and FPG SNPs meant that 7 FPG loci were also in the list of type 2 diabetes loci. The median genotyping call rate in the DIP samples was 0.992 (range 0.981-0.996), and there was >99% concordance between duplicate samples (8% of total genotyped samples were duplicates). We excluded one FPG SNP and one type 2 diabetes SNP that showed deviation from Hardy-Weinberg Equilibrium (Bonferroni-corrected *P* value <0.05). For details of included and excluded SNPs and their sources, see Table 1.

**Table 1a.**
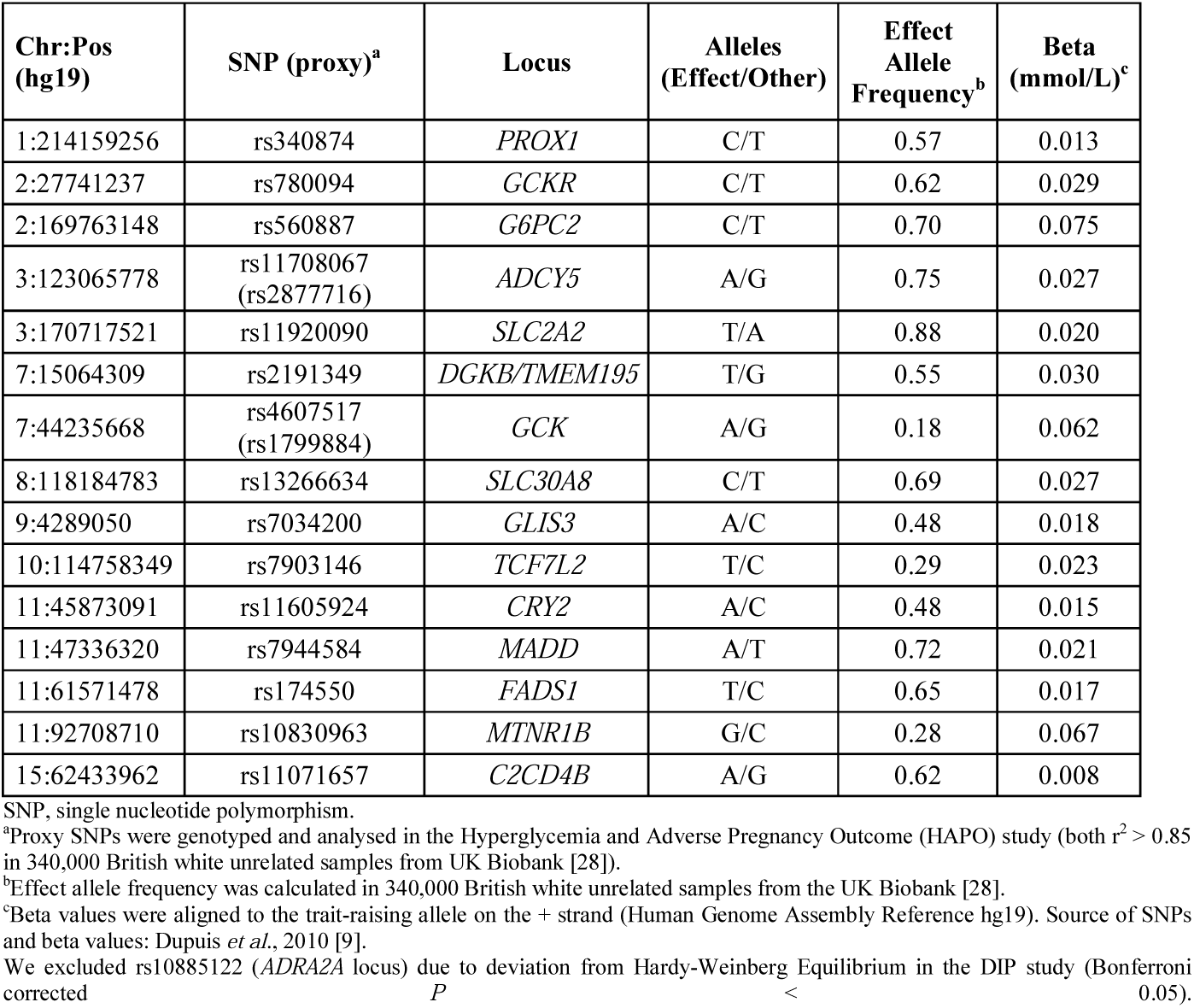
Fifteen SNPs associated with fasting plasma glucose (FPG) and used to construct the FPG genetic score.

**Table 1b.** Thirty-eight SNPs associated with type 2 diabetes (T2D) risk and used to construct the T2D genetic score.

| **Chr:Pos**<br>**(hg19)** | **SNP**<br>**(proxy)<sup>a</sup>** | **Locus** | **Alleles**<br>**(Effect/Other)** | **Effect**<br>**Allele**<br>**Frequency<sup>b</sup>** | **Beta<sup>c</sup>** | **Source in**<br>**which SNP**<br>**was**<br>**originally**<br>**identified** |
| --- | --- | --- | --- | --- | --- | --- |
| 1:120526982 | rs1493694 | *NOTCH2* | T/C | 0.11 | 0.110 | Zeggini *et al.*,<br>2008 [29] |
| 1:214163675 | rs340835 | *PROX1* | A/G | 0.49 | 0.062 | Dupuis *et al.*,<br>2010 [9] |
| 2:27741237 | rs780094 | *GCKR* | C/T | 0.62 | 0.011 | Dupuis *et al.*,<br>2010 [9] |
| 2:43732823 | rs7578597 | *THADA* | T/C | 0.89 | 0.141 | Zeggini *et al.*,<br>2008 [29] |
| 2:60584819 | rs243021 | *BCL11A* | A/G | 0.46 | 0.090 | Voight *et al.*,<br>2010 [30] |
| 2:227093745 | rs2943641<br>(rs2943640) | *IRS1* | C/T | 0.65 | 0.083 | Rung *et al.*,<br>2009 [31] |
| 3:12393125 | rs1801282 | *PPARG* | C/G | 0.88 | 0.138 | Altshuler *et al.*, 2000 [32] |
| 3:23336450 | rs7612463 | *UBE2E2* | C/A | 0.89 | 0.102 | Yamauchi *et al.*, 2010 [33] |
| 3:64711904 | rs4607103<br>(rs6795735) | *ADAMTS9* | C/T | 0.76 | 0.092 | Zeggini *et al.*,<br>2008 [29] |
| 3:123065778 | rs11708067<br>(rs2877716) | *ADCY5* | A/G | 0.75 | 0.097 | Dupuis *et al.*,<br>2010 [9] |
| 4:6292915 | rs10010131 | *WFS1* | G/A | 0.60 | 0.104 | Sandhu *et al.*,<br>2007 [34] |
| 5:76424949 | rs4457053 | *ZBED3* | G/A | 0.32 | 0.150 | Voight *et al.*,<br>2010 [30] |
| 6:20661250 | rs7754840<br>(rs9368222) | *CDKAL1* | C/G | 0.31 | 0.170 | Zeggini *et al.*,<br>2007 [35] |
| 7:28189411 | rs1635852 | *JAZF1* | T/C | 0.49 | 0.120 | Zeggini *et al.*,<br>2008 [29] |
| 7:44235668 | rs4607517<br>(rs1799884) | *GCK* | A/G | 0.18 | 0.029 | Dupuis *et al.*,<br>2010 [9] |
| 7:130466854 | rs972283<br>(rs4731702) | *KLF14* | G/A | 0.51 | 0.099 | Kong *et al.*,<br>2009 [36] |
| 8:95937502 | rs7845219 | *TP53INP1* | T/C | 0.50 | 0.093 | Voight *et al.*,<br>2010 [4] |
| 8:118184783 | rs13266634 | *SLC30A8* | C/T | 0.69 | 0.139 | Sladek *et al.*,<br>2007 [37] |
| 9:22133284 | rs10965250 | *CDKN2A/B* | G/A | 0.83 | 0.181 | Zeggini *et al.*,<br>2007 [35] |
| 9:81952128 | rs13292136 | *CHCHD9* | C/T | 0.93 | 0.182 | Voight *et al.*,<br>2010 [30] |

| Chr:Pos<br>(hg19) | SNP<br>(proxy)<sup>a</sup> | Locus | Alleles<br>(Effect/Other) | Effect<br>Allele<br>Frequency<sup>b</sup> | Beta<sup>c</sup> | Source in<br>which SNP<br>was<br>originally<br>identified |
| --- | --- | --- | --- | --- | --- | --- |
| 10:12328010 | rs12779790<br>(rs11257655) | *CDC123/*<br>*CAMK1D* | G/A | 0.18 | 0.088 | Zeggini *et al.*,<br>2008 [29] |
| 10:94465559 | rs5015480 | *HHEX/IDE* | C/T | 0.59 | 0.166 | Zeggini *et al.*,<br>2007 [35] |
| 10:114758349 | rs7903146 | *TCF7L2* | T/C | 0.29 | 0.335 | Grant *et al.*,<br>2006 [38] |
| 11:1696849 | rs2334499 | *HCCA2/DUSP8* | T/C | 0.42 | 0.080 | Kong *et al.*,<br>2009 [36] |
| 11:2691471 | rs231362 | *KCNQ1* | G/A | 0.52 | 0.104 | Kong *et al.*,<br>2009 [36] |
| 11:2847069 | rs163184 | *KCNQ1* | G/T | 0.48 | 0.083 | Yasuda *et al.*,<br>2008 [39],<br>Unoki *et al.*,<br>2008 [40] |
| 11:17408630 | rs5215 | *KCNJ11* | C/T | 0.36 | 0.089 | Gloyn *et al.*,<br>2003 [41] |
| 11:72433098 | rs1552224 | *CENTD2* | A/C | 0.84 | 0.123 | Voight *et al.*,<br>2010 [30] |
| 11:92673828 | rs1387153<br>(rs10830963) | *MTNR1B* | T/C | 0.29 | 0.115 | Prokopenko<br>*et al.*, 2009<br>[42], Dupuis<br>*et al.*, 2010<br>[9] |
| 12:66170163 | rs2612067 | *HMGA2* | G/T | 0.10 | 0.180 | Voight *et al.*,<br>2010 [30] |
| 12:71613276 | rs1353362 | *TSPAN8/ LGR5* | C/T | 0.28 | 0.103 | Zeggini *et al.*,<br>2008 [29] |
| 12:121402932 | rs7305618<br>(rs12427353) | *HNF1A* | C/T | 0.77 | 0.112 | Voight *et al.*,<br>2010 [30] |
| 13:80717156 | rs1359790 | *SPRY2* | G/A | 0.71 | 0.096 | Shu *et al.*,<br>2010 [43] |
| 15:62396389 | rs7172432 | *C2CD4A/B* | A/G | 0.57 | 0.068 | Yamauchi *et al.*, 2010 [33] |
| 15:77747190 | rs7178572 | *HMG20A* | G/A | 0.71 | 0.068 | Kooner *et al.*,<br>2011 [44] |
| 15:80432222 | rs11634397 | *ZFAND6* | G/A | 0.66 | 0.102 | Voight *et al.*,<br>2010 [30] |
| 17:36098040 | rs4430796 | *HNF1B* | G/A | 0.48 | 0.130 | Gudmundsson<br>*et al.*, 2007<br>[45] |
| X:152908152 | rs2301142 | *DUSP9* | A/G | 0.85 | 0.086 | Voight *et al.*,<br>2010 [30] |

SNP, single nucleotide polymorphism.
<sup>a</sup>Proxy SNPs were genotyped and analysed in the Hyperglycemia and Adverse Pregnancy Outcome (HAPO) study ( $r^2 > 0.7$ in 340,000 British white unrelated samples from UK Biobank [28], except for at *ADAMTS9* where $r^2 = 0.45$ [28]).
<sup>b</sup>Effect allele frequency was calculated in 340,000 British white unrelated samples from the UK Biobank [28].
<sup>c</sup>Beta values were aligned to the T2D-risk allele on the + strand (Human Genome Assembly Reference hg19). Beta value = log odds ratio for T2D from genome-wide association study meta-analysis of up to 8130 cases and 38987 controls, published in Voight *et al.* 2010 [30].
We excluded rs8042680 (*PRCI* locus, DIP study) and rs1470579 (*IGF2BP2* locus, HAPO study) from the T2D GS due to deviation from Hardy-Weinberg Equilibrium (Bonferroni-corrected $P < 0.05$ ). We additionally excluded rs11642841 (*FTO* locus) due to its primary effect on BMI [22].

In the HAPO study, we selected SNPs from the same 16 FPG and 41 type 2 diabetes loci for genotyping in women of European ancestry with DNA available. The selection and genotyping of SNPs in the HAPO study was performed at different times from that in the DIP study. Owing to the differing availability of published GWAS results at these times, the genotyped SNPs differed between HAPO and DIP at 9 of the associated loci. The HAPO SNPs at the 9 loci were generally well correlated with those genotyped in DIP (r^2^ >0.7, apart from at the *ADAMTS9* locus where r^2^ = 0.45). The median genotyping call rate in the HAPO samples was 0.984 (range 0.955-0.991), and the mean concordance between duplicate samples was >98.5% (at least 1% of samples were duplicated). We excluded 1 SNP that showed deviation from Hardy-Weinberg Equilibrium in the HAPO study (Bonferroni-corrected *P* value <0.05; see Table 1). After exclusion of SNPs that showed deviation from Hardy-Weinberg equilibrium and one SNP from the type 2 diabetes score whose main effect was on BMI (rs11642841 (*FTO* locus) [22]), a total of 15 SNPs at FPG-associated loci and 38 SNPs at type 2 diabetes-associated loci were available in both studies for analysis.

### Generating a genetic score for FPG and type 2 diabetes

Weighted genetic scores for FPG (FPG GS) and type 2 diabetes (T2D GS) were generated using the 15 SNPs and 38 SNPs, respectively. The GSs were calculated by taking the sum of the number of FPG-raising or type 2 diabetes risk alleles (0, 1 or 2) for each SNP, multiplied by its corresponding beta value (effect size) for association with FPG or type 2 diabetes, divided by the sum of all beta values and multiplied by the total number of SNPs analyzed (see Figure 2 for formula). GS were generated for participants with complete data for all included SNPs only.

**Figure 2.**
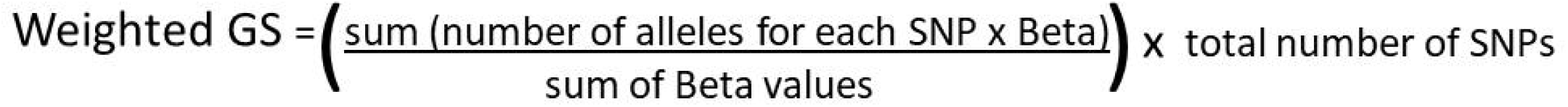
Formula for generating a weighted genetic score (GS). “Number of alleles” corresponds to either the number of risk alleles (T2D SNPs) or the number of glucose-raising alleles (FPG SNPs). FPG, fasting plasma glucose; SNP, single-nucleotide polymorphism; T2D, type 2 diabetes.

### Statistical analyses

#### Analysis of clinical characteristics

Clinical characteristics were compared between participants with and without GDM in HAPO and DIP using unpaired *t*-tests for normally distributed data and the Wilcoxon Rank-Sum test for non-normally distributed data. *P* values were corrected for 24 comparisons using the Bonferroni method.

#### Analysis of associations between FPG GS or T2D GS with glucose levels and GDM

Associations of the FPG GS or T2D GS with FPG, 1-hour and 2-hour glucose in women with and without GDM (cases and controls) were analyzed using linear regression in HAPO (which was a representative sample of European participants from the whole study cohort) and *P* values corrected for 12 comparisons using the Bonferroni method. Means for FPG GS and T2D GS in women with and without GDM were compared using unpaired *t*-tests in each study cohort separately, as the genetic scores were higher overall in DIP. *P* values were Bonferroni corrected for 16 comparisons.

#### Statistical software

All statistical analyses were performed using Stata version 14.0 (StataCorp LP, College Station, TX, USA). *P* values <0.05 were considered to indicate evidence of association, unless otherwise stated.

### Ethics approval

Ethics approval was obtained from the Northwestern University Office for the Protection of Research Participants for HAPO. The HAPO study protocol was approved by the institutional review board at each field center and all participants gave written, informed consent. Ethics approval was obtained from the local Galway University Hospital Research Ethics Committee for Atlantic DIP and all participants gave written, informed consent.

## RESULTS

### Clinical characteristics in women with and without GDM

Clinical characteristics for women with and without GDM are summarized in Tables 2a and 2b for HAPO and DIP, respectively. Women with a FPG ≥5.1 mmol/L (on its own or with either 1-hour or 2-hour hyperglycemia) had a higher pre-pregnancy BMI than women without GDM in HAPO and DIP (*P* values <0.001). Women with both fasting and either 1-hour or 2-hour hyperglycemia were older compared with controls in HAPO (*P* value <0.05 after Bonferroni correction). In HAPO we observed a higher SBP for women diagnosed with GDM by a FPG ≥5.1 mmol/L only compared with controls (*P* value <0.001) and they had a higher SBP when either their 1-hour or 2-hour glucose was also raised, but the *P* value was >0.05 after Bonferroni correction. In DIP there was a higher SBP for women diagnosed by both fasting and either 1-hour or 2-hour hyperglycemia criteria compared with controls (*P* value <0.05 after Bonferroni correction).

**Table 2a.** Clinical characteristics for participants diagnosed with GDM by the different criteria in HAPO.

| **HAPO** | **Controls**<br>**with normal**<br>**glucose** | **FPG $\geq 5.1$**<br>**mmol/L**<br>**only** | **1-hr**<br>**glucose<sup>a</sup>**<br>**$\geq 10$**<br>**mmol/L**<br>**only** | **2-hr**<br>**glucose<sup>a</sup>**<br>**$\geq 8.5$**<br>**mmol/L**<br>**only** | **Both (FPG**<br>**$\geq 5.1$ mmol/L**<br>**and either 1-**<br>**hr glucose<sup>a</sup>**<br>**$\geq 10$ mmol/L**<br>**or 2-hr**<br>**glucose<sup>a</sup>$\geq 8.5$**<br>**mmol/L)** |
| --- | --- | --- | --- | --- | --- |
| **Median FPG**<br>**in mmol/L**<br>**(IQR)** | 4.5<br>(4.3-4.7)<br>n=2,275 | 5.2<br>(5.1-5.3)<br>n=164 | 4.8<br>(4.6-4.9)<br>n=66 | 4.5<br>(4.3-4.7)<br>n=48 | 5.3<br>(5.2-5.5)<br>n=75 |
| **Median 1-hr**<br>**glucose in**<br>**mmol/L**<br>**(IQR)** | 7.1<br>(6.0-8.0)<br>n=2,275 | 8.4<br>(7.6-9.2)<br>n=164 | 10.4<br>(10.2-11.0)<br>n=66 | 9.0<br>(8.6-9.5)<br>n=48 | 10.6<br>(10.0-11.2)<br>n=75 |
| **Median 2-hr**<br>**glucose in**<br>**mmol/L**<br>**(IQR)** | 5.8<br>(5.1-6.5)<br>n=2,275 | 6.6<br>(6.0-7.1)<br>n=164 | 7.4<br>(6.6-7.9)<br>n=66 | 8.9<br>(8.6-9.1)<br>n=48 | 7.9<br>(7.1-8.9)<br>n=75 |
| **Median**<br>**maternal age**<br>**in years**<br>**(IQR)** | 31<br>(26-34)<br>n=2,275 | 31<br>(27-35)<br>n=164 | 31<br>(27-35)<br>n=66 | 32<br>(27-34)<br>n=48 | 32<br>(29-36)**<br>n=75 |
| **Median pre-**<br>**pregnancy**<br>**BMI (IQR)** | 22.9<br>(21.0-26.1)<br>n=2,125 | 27.5<br>(23.8-33.1)***<br>n=142 | 24.4<br>(21.2-27.9)<br>n=59 | 23.0<br>(20.1-25.1)<br>n=45 | 28.0<br>(23.8-35.2)***<br>n=65 |
| **Median SBP**<br>**in mmHg**<br>**(IQR)** | 108<br>(102-114)<br>n=2,275 | 113<br>(106-119)***<br>n=164 | 110<br>(103-118)<br>n=66 | 104<br>(100-116)<br>n=48 | 110<br>(103-118)*<br>n=75 |

BMI, body mass index; FPG, fasting plasma glucose; GDM, gestational diabetes; HAPO, Hyperglycemia and Pregnancy Outcome Study; SBP, systolic blood pressure.
<sup>a</sup>The 1-hr and 2-hr glucose measures refer to the glucose level measured at 1 and 2 hours, respectively, following a 75 g oral glucose load as part of an oral glucose tolerance test.
\* $P$ value $<0.05$ for comparison with controls ( $>0.05$ after Bonferroni correction).
\*\* $P$ value $<0.001$ for comparison with controls ( $<0.05$ after Bonferroni correction).
\*\*\* $P$ value $<0.001$ for comparison with controls (remained $<0.001$ after Bonferroni correction).

**Table 2b.** Clinical characteristics for women diagnosed with GDM by the different criteria in DIP.

| **DIP** | **Controls**<br>**with normal**<br>**glucose** | **FPG $\geq 5.1$**<br>**mmol/L**<br>**only** | **1-hr**<br>**glucose<sup>a</sup>**<br>**$\geq 10$**<br>**mmol/L**<br>**only** | **2-hr**<br>**glucose<sup>a</sup>**<br>**$\geq 8.5$**<br>**mmol/L**<br>**only** | **Both (FPG**<br>**$\geq 5.1$ mmol/L**<br>**and either 1-**<br>**hr glucose<sup>a</sup>**<br>**or 2-hr**<br>**glucose<sup>a</sup> $\geq 8.5$**<br>**mmol/L)** |
| --- | --- | --- | --- | --- | --- |
| **Median FPG**<br>**in mmol/L**<br>**(IQR)** | 4.3<br>(4.1-4.5)<br>n=816 | 5.3<br>(5.2-5.5)<br>n=58 | 4.6<br>(4.4-4.8)<br>n=88 | 4.5<br>(4.2-4.7)<br>n=25 | 5.5<br>(5.2-5.9)<br>n=97 |
| **Median 1-hr**<br>**glucose<sup>a</sup> in**<br>**mmol/L**<br>**(IQR)** | 6.6<br>(5.6-7.7)<br>n=816 | 8.7<br>(7.5-9.1)<br>n=58 | 10.8<br>(10.2-11.2)<br>n=88 | 8.6<br>(8.1-9.1)<br>n=25 | 11.2<br>(10.2-12.0)<br>n=97 |
| **Median 2-hr**<br>**glucose<sup>a</sup> in**<br>**mmol/L**<br>**(IQR)** | 5.2<br>(4.6-6.0)<br>n=816 | 6.1<br>(5.5-7.0)<br>n=58 | 6.9<br>(5.9-7.8)<br>n=88 | 8.8<br>(8.6-9.2)<br>n=25 | 8.5<br>(7.5-9.3)<br>n=97 |
| **Median**<br>**maternal age**<br>**in years**<br>**(IQR)** | 32<br>(29-36)<br>n=521 | 35<br>(31-39)*<br>n=35 | 34<br>(31-37)*<br>n=69 | 32<br>(29-40)<br>n=16 | 33<br>(30-36)<br>n=72 |
| **Median pre-**<br>**pregnancy**<br>**BMI (IQR)** | 25.4<br>(23.4-28.8)<br>n=454 | 31.6<br>(29.0-38.3)***<br>n=33 | 29.6<br>(25.5-35.7)***<br>n=56 | 28.5<br>(25.5-31.1)<br>n=16 | 33.5<br>(28.3-37.6)***<br>n=55 |
| **Median SBP**<br>**in mmHg**<br>**(IQR)** | 117<br>(108-124)<br>n=437 | 119<br>(110-130)<br>n=21 | 120<br>(113-130)*<br>n=38 | 122<br>(111-134)<br>n=12 | 120<br>(115-134)**<br>n=41 |

BMI, body mass index; DIP, Atlantic Diabetes in Pregnancy; FPG, fasting plasma glucose; GDM, gestational diabetes; SBP, systolic blood pressure.
<sup>a</sup>The 1-hr and 2-hr glucose measures refer to the glucose level measured at 1 and 2 hours, respectively, following a 75 g oral glucose load as part of an oral glucose tolerance test.
\* $P$ value $<0.01$ for comparison with controls ( $>0.05$ after Bonferroni correction).
\*\* $P$ value $<0.01$ for comparison with controls ( $<0.05$ after Bonferroni correction).
\*\*\* $P$ value $<0.001$ for comparison with controls (remained $<0.001$ after Bonferroni correction).

**Table 3.**
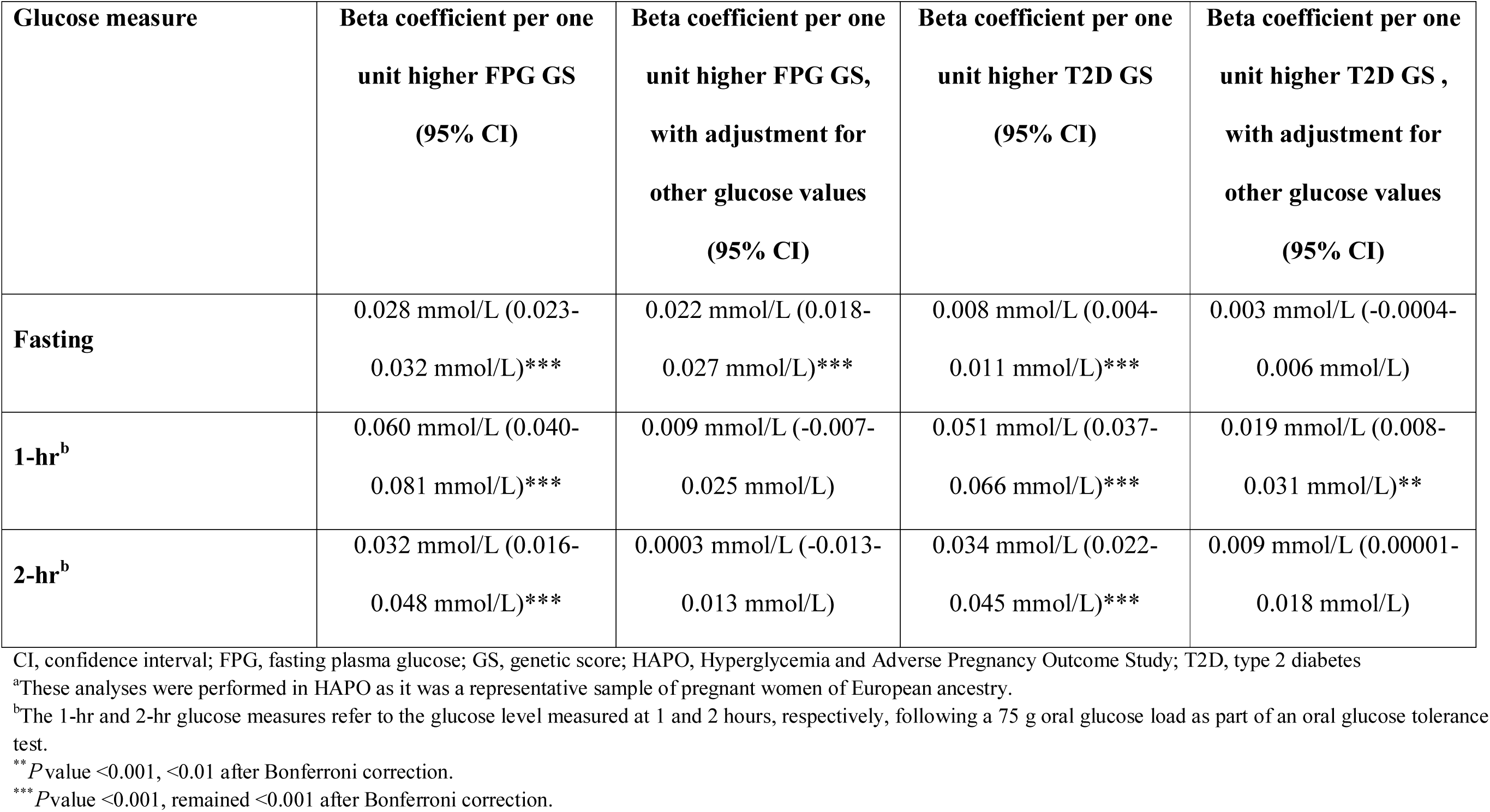
Associations for FPG and T2D GS with different measures of glucose tolerance in women with and without diabetes in HAPO^a^.

| **Glucose measure** | **Beta coefficient per one**<br>**unit higher FPG GS**<br>**(95% CI)** | **Beta coefficient per one**<br>**unit higher FPG GS,**<br>**with adjustment for**<br>**other glucose values**<br>**(95% CI)** | **Beta coefficient per one**<br>**unit higher T2D GS**<br>**(95% CI)** | **Beta coefficient per one**<br>**unit higher T2D GS ,**<br>**with adjustment for**<br>**other glucose values**<br>**(95% CI)** |
| --- | --- | --- | --- | --- |
| **Fasting** | 0.028 mmol/L (0.023-0.032 mmol/L)*** | 0.022 mmol/L (0.018-0.027 mmol/L)*** | 0.008 mmol/L (0.004-0.011 mmol/L)*** | 0.003 mmol/L (-0.0004-0.006 mmol/L) |
| **1-hr<sup>b</sup>** | 0.060 mmol/L (0.040-0.081 mmol/L)*** | 0.009 mmol/L (-0.007-0.025 mmol/L) | 0.051 mmol/L (0.037-0.066 mmol/L)*** | 0.019 mmol/L (0.008-0.031 mmol/L)** |
| **2-hr<sup>b</sup>** | 0.032 mmol/L (0.016-0.048 mmol/L)*** | 0.0003 mmol/L (-0.013-0.013 mmol/L) | 0.034 mmol/L (0.022-0.045 mmol/L)*** | 0.009 mmol/L (0.00001-0.018 mmol/L) |

CI, confidence interval; FPG, fasting plasma glucose; GS, genetic score; HAPO, Hyperglycemia and Adverse Pregnancy Outcome Study; T2D, type 2 diabetes
<sup>a</sup>These analyses were performed in HAPO as it was a representative sample of pregnant women of European ancestry.
<sup>b</sup>The 1-hr and 2-hr glucose measures refer to the glucose level measured at 1 and 2 hours, respectively, following a 75 g oral glucose load as part of an oral glucose tolerance test.
\*\**P* value <0.001, <0.01 after Bonferroni correction.
\*\*\**P* value <0.001, remained <0.001 after Bonferroni correction.

### FPG, 1-hour and 2-hour glucose are associated with FPG and T2D GS in pregnant women with and without GDM

FPG, 1-hour and 2-hour glucose values were associated with the fasting and type 2 diabetes genetic scores in HAPO (Table 2). Adjusting for the different measures of glucose tolerance suggested that these associations were not independent of one another.

### Women diagnosed with GDM by fasting glucose criteria have a higher FPG GS

We observed a higher FPG GS in women diagnosed with GDM by fasting hyperglycemia only and by both fasting and either 1-hour or 2-hour criteria, compared with controls (Figure 3A, all *P* values for comparison with control group <0.05 after Bonferroni correction). There was also evidence that women with a raised 1-hour glucose only had a higher FPG GS in HAPO (*P* value for comparison with controls <0.01 but >0.05 with Bonferroni correction), but this was not as strong in DIP (*P* value =0.05). In contrast, women diagnosed with GDM by 2-hour only criteria did not have a higher FPG GS overall (*P* values for comparison with controls >0.05 in both studies).

**Figure 3.**
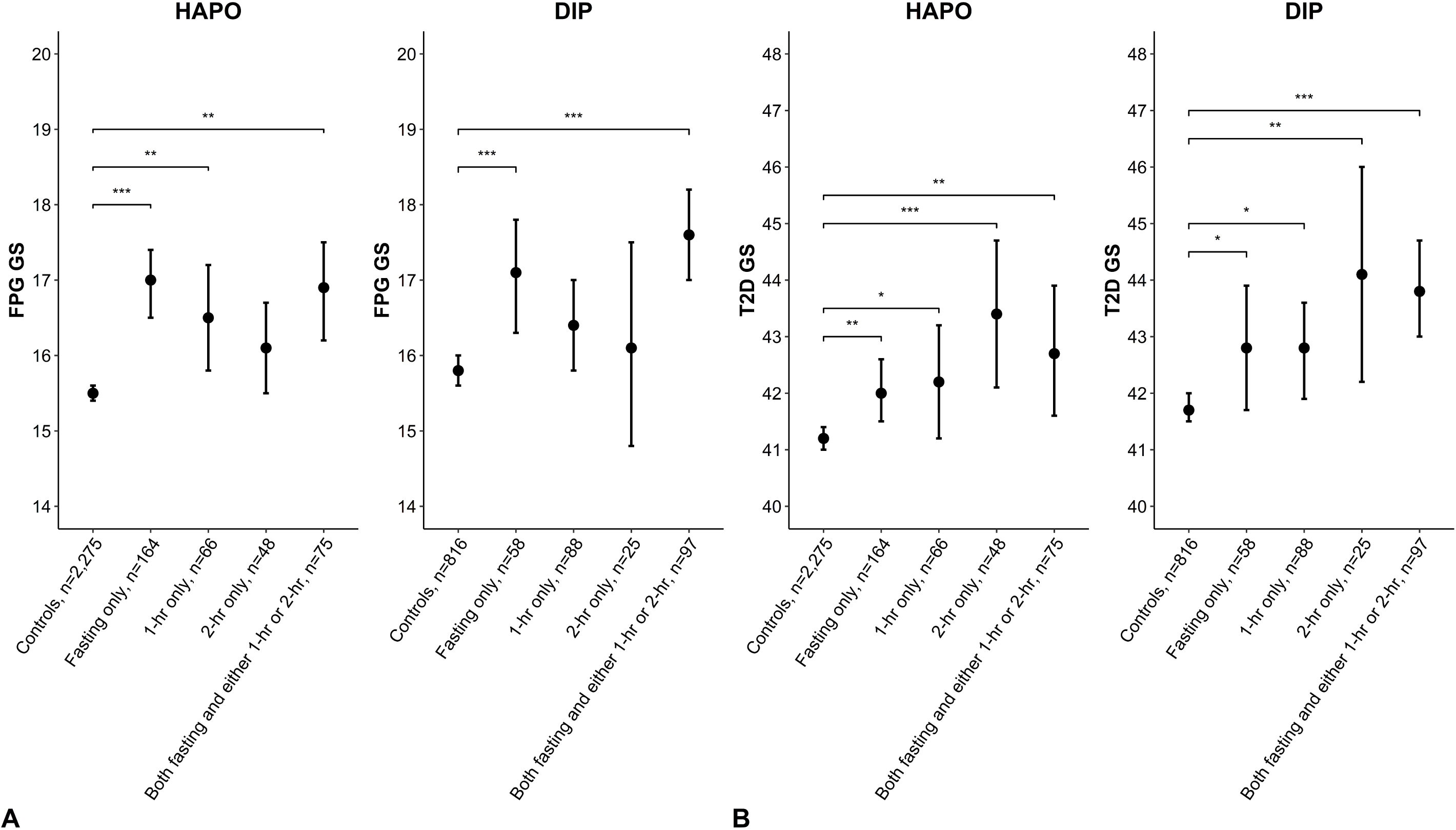
Plots showing mean FPG GS (A) or T2D GS (B) in each GDM glucose diagnostic category in HAPO and DIP. The 1-hr and 2-hr glucose groups refer to glucose levels measured at 1 and 2 hours, respectively, following a 75 g oral glucose load as part of an oral glucose tolerance test. The control group include women with a FPG <5.1 mmol/L, 1-hr glucose <10 mmol/L and 2-hour glucose <8.5 mmol/L. The fasting only group includes women with a FPG ≥5.1 mmol/L, a 1-hr glucose <10 mmol/L and 2-hour glucose <8.5 mmol/L. The 1-hr only group includes women with 1-hr glucose ≥10 mmol/L, FPG <5.1 mmol/L and 2-hr glucose <8.5 mmol/L. The 2-hr only group includes women with a 2-hr glucose ≥8.5 mmol/L, FPG <5.1 mmol/L and 1-hr glucose <10 mmol/L. The remaining group includes women with both a FPG ≥5.1 mmol/L and either a 1-hr glucose ≥10mmol/L or 2-hr glucose ≥8.5 mmol/L, or both. Error bars show 95% confidence intervals. \**P* value for comparison between cases and controls <0.05 \*\**P* value for comparison between cases and controls <0.01. \*\*\**P* value for comparison between cases and controls <0.001. All *P* values survived Bonferroni correction at α=0.05 except for the FPG GS in women with 1-hour hyperglycemia in HAPO and the T2D GS in women with isolated fasting or 1-hour hyperglycemia in HAPO and DIP. DIP; Atlantic Diabetes in Pregnancy; GDM, gestational diabetes; FPG GS, fasting plasma glucose genetic score; HAPO; Hyperglycemia and Adverse Pregnancy Outcome Study; OR, odds ratio; T2D GS, type 2 diabetes genetic score.

### Women diagnosed with GDM by fasting, 1-hour or 2-hour criteria have a higher T2D GS than controls

The T2D GS was higher than controls in women with fasting, 1-hour or 2-hour hyperglycemia in HAPO and DIP (Figure 2B): all *P* values for comparison with controls were <0.05 after correction except for the fasting and 1-hour only groups.

## CONCLUSIONS

In this study of 3,712 pregnant women of European ancestry, we have confirmed that women diagnosed with GDM according to the WHO 2013 criteria have a raised genetic risk for type 2 diabetes and shown for the first time that this risk was raised across all of the different measures of glucose tolerance. A genetic predisposition to a higher FPG was present for women who met the fasting glucose criteria (and 1-hour glucose criteria in HAPO), but was not present for women who met the 2-hour criteria.

We confirmed that FPG in pregnant women both with and without GDM was positively associated with a FPG GS generated using SNPs identified in a non-pregnant population [9]. The 1-hour and 2-hour glucose values were also correlated with the FPG GS, but this could potentially be explained by their association with FPG, since this association was not as strong once this was taken into account. Thus, the observation that the FPG GS was not higher in women diagnosed with GDM due to a 2-hour glucose ≥8.5 mmol/L alone was expected. Maternal FPG was also associated with the T2D GS, which would be expected, as there are loci within the T2D GS which also raise fasting glucose (e.g. *GCK, MTNR1B*) [9]. The *ADCY5* locus has also been found to be associated with 2-hour glucose values [23]. Thus, the observation of a higher T2D GS in women meeting the fasting or 2-hour WHO 2013 criteria for GDM is not surprising. A GWAS for 1-hour glucose values was not available at the time of writing, but since we found the T2D GS to be associated with 1-hour glucose values in HAPO, it is likely that this explains the higher T2D GS seen in the women meeting this criterion for diagnosis of GDM, and will contribute to the higher T2D GS seen in women with both a fasting and either 1-hour or 2-hour hyperglycaemia. However, it is important to note that the relationships between the T2D GS and the different glucose categories did not appear to be independent of one another, and again, although women meeting the diagnosis for GDM in one category may not meet the thresholds for GDM in other categories, they are likely to have a degree of fasting and postprandial hyperglycemia which will contribute to their higher genetic risk for type 2 diabetes compared with women without GDM.

One might expect that women with both fasting and postprandial hyperglycemia would have the highest genetic risk for type 2 diabetes, but we did not observe this for the T2D GS in women with both a FPG ≥5.1 mmol/L and either a 1-hour glucose ≥10 mmol/L or 2-hour glucose ≥8.5 mmol/L. On the whole, the relationship between GDM and a higher T2D GS was clearest for women with a raised 2-hour glucose or a combination of raised fasting and 1-hour or 2-hour glucose, but studies with greater statistical power will be needed to confirm whether genetic risk of T2D is heterogeneous across the different thresholds of glucose tolerance that are part of the WHO 2013 criteria for GDM.

This work specifically examining the genetic risk of type 2 diabetes in women diagnosed with GDM according to different measures of glucose tolerance supports the results from the recent HAPO Follow-Up Study [24] which showed that women diagnosed with GDM post-hoc according to WHO 2013 criteria had a higher risk for type 2 diabetes 10 to 14 years after pregnancy. We observed the highest BMIs in women diagnosed with GDM by fasting hyperglycemia only or both criteria, which is consistent with previous research showing that women diagnosed with GDM by the WHO 2013 criteria were more overweight than those diagnosed by WHO 1996 criteria [7,25]. However, the associations seen for GDM with FPG GS and T2D GS are not driven by BMI (the genetic variants included within the scores do not primarily affect FPG and T2D risk because of an effect on BMI), suggesting that women with fasting hyperglycemia in pregnancy are likely to have both BMI-related metabolic factors and a genetic predisposition contributing to type 2 diabetes risk. In the longer-term, although using the lower FPG threshold for identifying GDM will result in more cases diagnosed, these women will be an important target for long-term follow-up. The Diabetes Prevention Program (DPP) [26] trial found that lifestyle intervention or metformin treatment reduced risk of progression to type 2 diabetes in women with impaired glucose tolerance and a history of GDM (according to relevant criteria at time of diagnosis), but a genetic risk score for type 2 diabetes did not influence treatment response [27]. It is not known whether this would be different for women specifically diagnosed by WHO 2013 criteria, but it is likely that these women would benefit from monitoring after pregnancy.

There are limitations of this study that are important to consider. The small number of cases of GDM included has been mentioned and could explain why there were not clear differences in T2D GS seen between the different diagnostic categories. We also studied women from two different studies, where there were notable differences in clinical characteristics, even for women without GDM. Additionally, the FPG and T2D GS were consistently higher in DIP than in HAPO. This is likely to reflect differences in SNPs used to generate the genetic scores and possibly a slightly higher genetic disposition to a raised FPG and type 2 diabetes in DIP. However, there were remarkably similar patterns for the genetic score associations amongst the different diagnostic groups in both studies. The results of these analyses are therefore likely to be applicable to women of European ancestry, but further larger-scale studies, including analysis of women with diverse ancestry, will be needed to confirm the associations identified in this study.

In conclusion, women diagnosed with GDM according to the newest WHO 2013 criteria, regardless of how the diagnosis is met, have a higher genetic risk for type 2 diabetes compared with women without GDM. Overall, the criteria identify an important group of women at risk for adverse pregnancy outcomes as well as a higher risk for developing future type 2 diabetes [8]. This study has added to the literature confirming genetic predisposition to type 2 diabetes in women with GDM and supports the possibility that genetic testing could be a novel tool to help identify women at high risk for GDM at an early stage of pregnancy, helping to target screening and early intervention.

## DATA AVAILABILITY

Data analysed and generated in this study are available to researchers through open collaboration. For access to the data used in this study please contact Dr Rachel Freathy and Professor William Lowe Jr in relation to HAPO and Dr Rachel Freathy and Professor Fidelma Dunne in relation to Atlantic DIP. The websites describing the studies and other data available are https://www.ncbi.nlm.nih.gov/projects/gap/cgibin/study.cgi?study_id=phs000096.v4.p1 for HAPO and http://atlanticdipireland.com/ for Atlantic DIP.

## AUTHOR CONTRIBUTIONS

AEH carried out analyses, wrote the manuscript, reviewed and edited the manuscript and contributed to the discussion. GMH was involved in the original HAPO analyses, reviewed and edited the manuscript and contributed to the discussion. AME was involved in the original DIP analyses, reviewed and edited the manuscript and contributed to the discussion. KAP researched data, reviewed and edited the manuscript and contributed to the discussion. DMS was involved in the original HAPO analyses, reviewed and edited the manuscript and contributed to the discussion. LPL led the collection and preparation of the HAPO samples for genotyping, reviewed and edited the manuscript and contributed to the discussion. WLL was involved in the original data acquisition and analysis in HAPO, reviewed and edited the manuscript and contributed to the discussion. FPD was involved in the original data acquisition and analysis in DIP, reviewed and edited the manuscript and contributed to the discussion. ATH researched data, reviewed and edited the manuscript and contributed to the discussion. RMF researched data, wrote the manuscript, reviewed and edited the manuscript and contributed to the discussion. WLL (HAPO), FDP (DIP) and RMF are the guarantors of this work, accept full responsibility for the work and/or the conduct of the study, had access to the data, and controlled the decision to publish.

## ACKNOWLEDGEMENTS

We acknowledge the work of the HAPO and DIP original investigators, whose names can be viewed in their original publications. We acknowledge the role of all professionals and families who contributed to HAPO and DIP.

## PRIOR PUBLICATION

Parts of this work were presented at the European Association for the Study of Diabetes Annual Meeting, Stockholm, Sweden, 14-18 September 2015, the Royal College of Obstetricians and Gynaecologists Annual Academic Meeting, 8-9 February 2018, the East of England Deanery Registrar Prize Meeting, 15 June 2018 and the European Association for the Study of Diabetes Diabetic Pregnancy Study Group Annual Meeting, Graz, Austria, 5-8 September 2019. A previous version of the manuscript was posted as a preprint on bioRxiv, 22 June 2020 (doi: https://doi.org/10.1101/671057).

